# Efficacy of Targeting SARS-CoV-2 by CAR-NK Cells

**DOI:** 10.1101/2020.08.11.247320

**Authors:** Minh Ma, Saiaditya Badeti, Ke Geng, Dongfang Liu

## Abstract

SARS-CoV-2, which causes COVID-19 disease, is one of greatest global pandemics in history. No effective treatment is currently available for severe COVID-19 disease. One strategy for implementing cell-based immunity involves the use of chimeric antigen receptor (CAR) technology. Unlike CAR T cells, which need to be developed using primary T cells derived from COVID-19 patients with lymphopenia, clinical success of CAR NK cell immunotherapy is possible through the development of allogeneic, universal, and ‘off-the-shelf’ CAR-NK cells from a third party, which will significantly broaden the application and reduce costs. Here, we develop a novel approach for the generation of CAR-NK cells for targeting SARS-CoV-2. CAR-NK cells were generated using the scFv domain of CR3022 (henceforth, CR3022-CAR-NK), a broadly neutralizing antibody for SARS-CoV-1 and SARS-CoV-2. CR3022-CAR-NK cells can specifically bind to RBD of SARS-CoV-2 and pseudotyped SARS-CoV-2 S protein, and can be activated by pseudotyped SARS-CoV-2-S viral particles *in vitro.* Further, CR3022-CAR-NK cells can specifically kill pseudo-SARS-CoV-2 infected target cells. Thus, ‘off-the-shelf’ CR3022-CAR-NK cells may have the potential to treat patients with severe COVID-19 disease.

## INTRODUCTION

SARS-CoV-2 is highly contagious, which presents a significant public health issue^1^. Currently, there is no vaccine available^2^. An FDA-approved standard of treatment for COVID-19 is not available either. Current treatment for COVID-19 patients can be classified into three categories: anti-viral treatments^3^, immunosuppression-based treatments^4^, and other supporting treatments such as convalescent plasma^5^. Specifically, in a few trials patients have been given combinations of antivirals including umifenovir^6^, remdesivir/ribavirin^7^, chloroquine^8^ (an anti-malarial drug), chloroquine’s analogue hydroxychloroquine^9^(a disease-modifying antirheumatic drug), and/or lopinavir/ritonavir^10,11^. Non-steroidal anti-inflammatory drugs (NSAIDs), antibodies against IL-6 receptors, and corticosteroids have also been used during the early acute phase of SARS-CoV-2 to suppress the overactivated immune response^10^. Other supporting therapies include supplemental oxygen and mechanical ventilatory support when indicated (e.g., intubation, etc.).

Given the recent success of immunotherapy in cancer^12^, several immune cell-based immunotherapeutic strategies against SARS-CoV-2 are being rapidly developed, which include quantification or adoptive transfer of monocytes or NK cells (<u>NCT04375176, NCT04280224</u>, and <u>NCT04365101</u>), universal ‘off-the-shelf’ NKG2D-ACE2 CAR-NK cells (<u>NCT04324996</u>), and several stem cell-based immunotherapeutic strategies (<u>NCT04416139</u>). Clinically, NK cells were first defined as CD56^bright^CD16^-^ and CD56^dim^CD16^+^ in the peripheral blood^13^. These NK cells isolated from peripheral blood can be further modified to express CAR for treating a variety of cancer and infectious diseases^14^.

In this study, we develop a novel approach for the generation of CAR-NK cells for targeting SARS-CoV-2. CAR-NK cells were generated using the scFv domain of CR3022, a strong neutralizing antibody for SARS-CoV-1 and SARS-CoV-2. The data show that ‘off-the-shel’ CR3022-CAR-NK cells may have the potential to treat patients with severe COVID-19 disease.

## RESULTS

### Generation of CR3022-CAR-NK-92MI cells

To develop an NK cell-based immunotherapy to treat COVID-19 patients, we modified NK cells with a CAR molecule specific against SARS-CoV-2 S protein. Previous studies show that the genome sequence of SARS-CoV-2 is highly similar to that of SARS-CoV-1^15^. Recent studies also demonstrate that a previously isolated neutralizing antibody from a convalescent SARS patient (and later named CR3022) can specifically bind to the RBD of SARS-CoV-2 spike protein^16^. Thus, we cloned the scFv domain of CR3022 into an SFG retroviral vector that contains a human IgG1 hinge and CH2-CH3 domain, CD28 transmembrane (TM) domain and intracellular domain, 4-1BB-Ligand intracellular domain, and CD3zeta intracellular domain (**Fig. 1a**). Specifically, we chose the scFv domain of CR3022 antibody because of its strong binding activity against both SARS-CoV-1 and SARS-CoV-2 S proteins.

**Figure 1:**
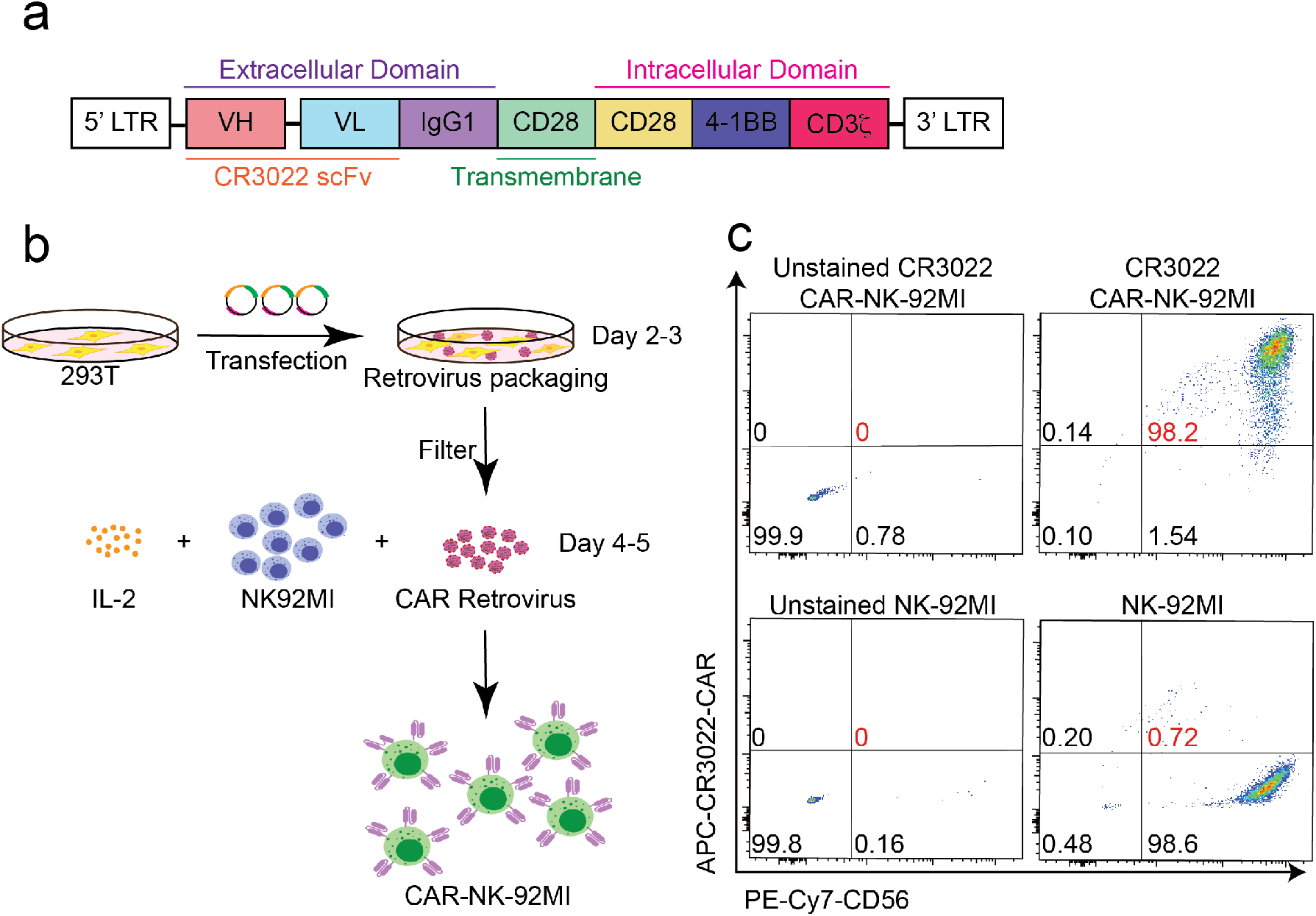
Generation of CR3022-CAR-NK92MI cells. (**a**) Schematic design of CR3022-CAR in SFG retroviral vector. The SFG retroviral vector contains the CR3022 single chain antibody fragment (clone 3), a human IgG1 CH2CH3 hinge region and CD28 transmembrane region, followed by the intracellular domains of co-stimulatory CD28, 4-1BB, and the intracellular domain of CD3ζ. (**b**) Generation of CR3022-CAR-NK cells. 293T cells were transfected with SFG-CR3022-CAR for 48-72 hours for CAR retrovirus packaging and transduced into NK92MI cells. (**c**) Determination of CAR expression by flow cytometry. CR3022-CAR cells were harvested after 4-5 days then stained with anti-CD56 and CAR F(ab)2 domain [IgG (H+L)] for flow cytometry.

To overcome the limitations of T cell therapy and take advantage of the benefits of using NK cells in targeting SARS-CoV-2 *in vivo,* we propose a CAR-expressing NK cell therapeutic approach. To initiate our studies, we successfully generated CR3022-CAR-NK cells in the human NK-92 cell line background (**Fig. 1a**). Specifically, the NK-92 cell line was transduced with CR3022-CAR (**Fig. 1**). Then, the subsequent CR3022-CAR positive NK-92 cells were sorted by flow cytometry. Sorted CR3022-CAR-NK-92MI cells were maintained for 2 months to verify CAR expression (data not shown). The generation of CR3022-CAR is schematically described (**Fig. 1b)**. 293T cells were transfected with a combination of plasmids containing CR3022-CAR in the SFG backbone, RDF, and PegPam3, as previously described^17,18^. The SFG retrovirus particles were used to transduce NK-92MI cells. After 4-5 days, NK-92MI and CR3022-CAR cells were stained with CD56 and human IgG (H+L) and the CAR expression was analyzed by flow cytometry (**Fig. 1c**). Greater than 98% of CD56^+^ CR3022-CAR^+^ NK-92MI cells were observed (**Fig. 1**). In summary, we have successfully established the stable membrane expression of CR3022-CAR-NK cells.

### Characteristics of CR3022-CAR-NK92MI cells

To have a better understanding of the immunophenotype of CR3022-CAR-NK-92MI, we further examined the expressions of several key immunoreceptors on CR3022-CAR-NK-92MI cells by flow cytometry. These receptors include TIGIT, LAG-3, TIM-3, KLRG1, CTLA-4, PD-1, CD69, CD8A, NKG2C, CD94, DNAM-1, 2B4, NKG2D, NKP46, and CD16 (**Fig. 2a**). The expressions of these activating and inhibitory receptors are comparable between parental NK-92MI and CR3022-CAR-NK-92MI cells. Surprisingly, the expressions of CD94 and 2B4 receptors significantly decreased in CR3022-CAR-NK-92MI cells. Overall, these key activating and inhibitory receptors between parental NK-92MI and CR3022-CAR-NK-92MI cells are similar, indicating the stable characteristics of NK-92MI at pre- and post-transduction stages.

**Figure 2:**
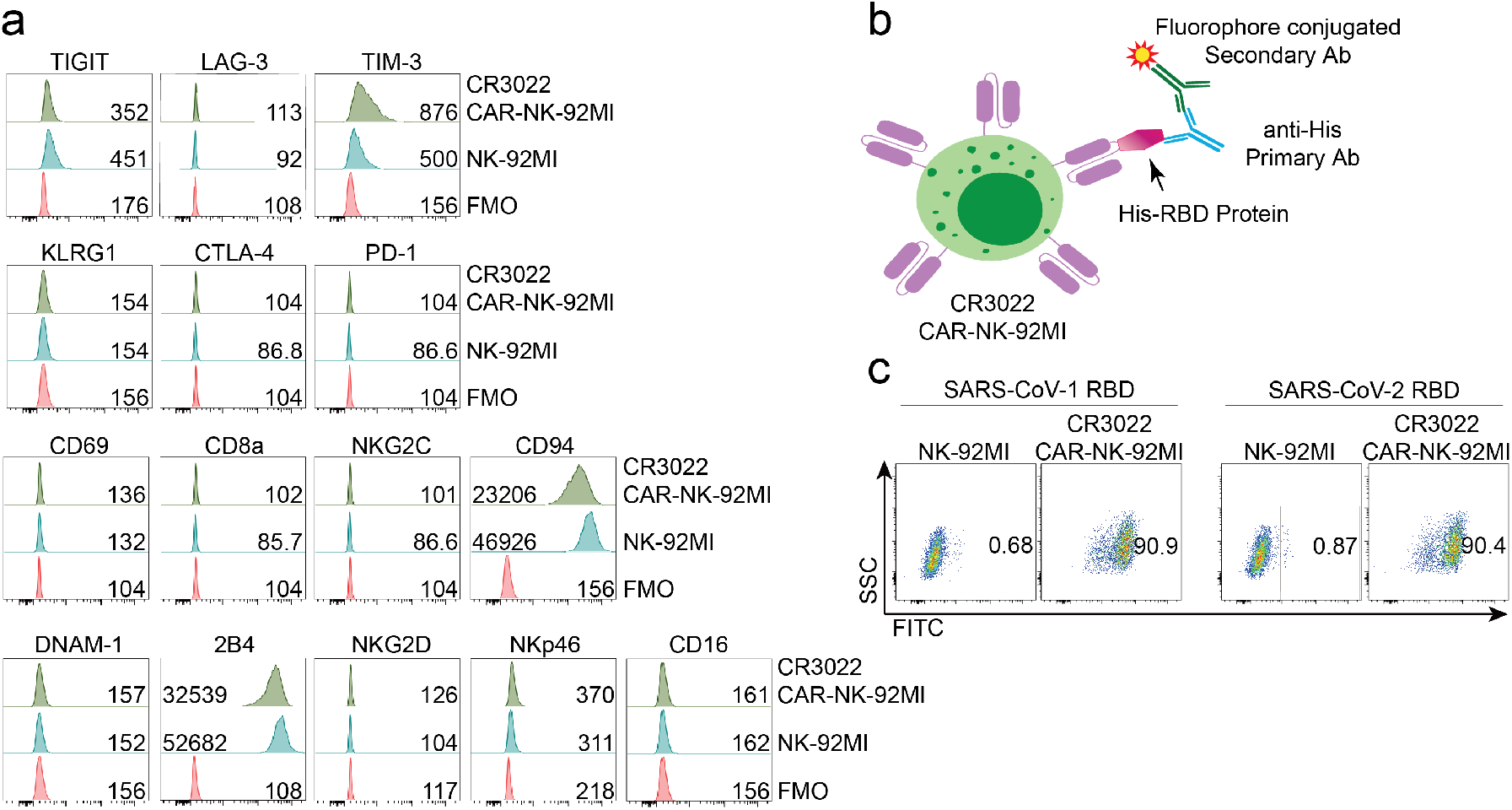
CR3022-CAR-NK92MI cells bind to RBD domain of SARS-CoV-2-S protein. (**a**) Immunophenotyping of CR3022-CAR. Antibodies against various immunomodulatory receptors including TIGIT, LAG-3, TIM-3, KLRG1, CTLA-4, PD-1, CD69, CD8A, NKG2C, CD94, DNAM-1, 2B4, NKG2D, NKP46, and CD16 were used to stain CR3022-CAR and NK-92MI. The expression of these receptors was determined by flow cytometry. (**b**) Diagram of CR3022-CAR binding to the RBD domain of SARS-CoV-2-S recombinant protein. CR3022-CAR binds to RBD of SARS-CoV-2-S protein which is then recognized by anti-His and its corresponding secondary antibody conjugated to a fluorophore. (**c**) Representative dot plots of CR3022-CAR binding to RBD of SARS-CoV-2. CR3022-CAR or NK-92MI cells were incubated with SARS-CoV-2-RBD or SARS-CoV-1-RBD recombinant protein. The binding efficiency was determined by flow cytometry.

After successful establishment of CR3022-CAR-NK-92MI cells, we then assessed the binding activity of CR3022-CAR cells to the RBD domain of SARS-CoV-2-S protein. CR3022-CAR-NK-92MI cells and NK-92MI cells were incubated with the recombinant His-RBD protein from SARS-CoV-1 or SARS-CoV-2, respectively. The complex of CR3022-CAR-NK-92MI and the His-RBD protein was then recognized by anti-His and its corresponding secondary antibody conjugated with a fluorophore (**Fig. 2b**). Flow cytometry was used to evaluate the binding efficiency of CR3022-CAR to the RBD of S protein from either SARS-CoV-1 or SARS-CoV-2. Consistent with earlier results from previous studies^16^, CR3022 binds to the RBD of both SARS-CoV-1 and SARS-CoV-2 (**Fig. 2c**). We therefore conclude that CR3022-CAR-NK-92MI cells can specifically bind to the recombinant His-RBD protein from SARS-CoV-1 and SARS-CoV-2.

### CR3022-CAR-NK cells bind to pseudotyped SARS-CoV-2-S viral particles

However, the partial RBD domain of SARS-CoV-2-S may not fully reflect the complexity of SARS-CoV-2 viral particles. Therefore, we evaluated the binding activity of CR3022-CAR cells to pseudotyped SARS-CoV-2-S viral particles purchased from GenScript, USA. Similar to the concept in **Figure 2b**, CR3022-CAR-NK-92MI binds to the pseudotyped SARS-CoV-2-S viral particles. The CR3022-CAR-NK-92MI and SARS-CoV-2-S viral particle complex can be recognized by the binding of anti-spike antibody and its corresponding fluorophore-conjugated secondary antibody (**Fig. 3a**). Previous studies show that the RBD of spike protein binds to ACE2 and facilitates SARS-CoV-2 entry^19^. As a positive control, we included 293T-hACE2 in our experiment (**Fig. 3b**). In addition, we also included spike recombinant proteins, full-length and S1 subunit containing RBD as an additional control group. As expected, CR3022-CAR-NK-92MI cells was able to bind to the pseudotyped SARS-CoV-2-S viral particles (**Fig. 3c**).

**Figure 3:**
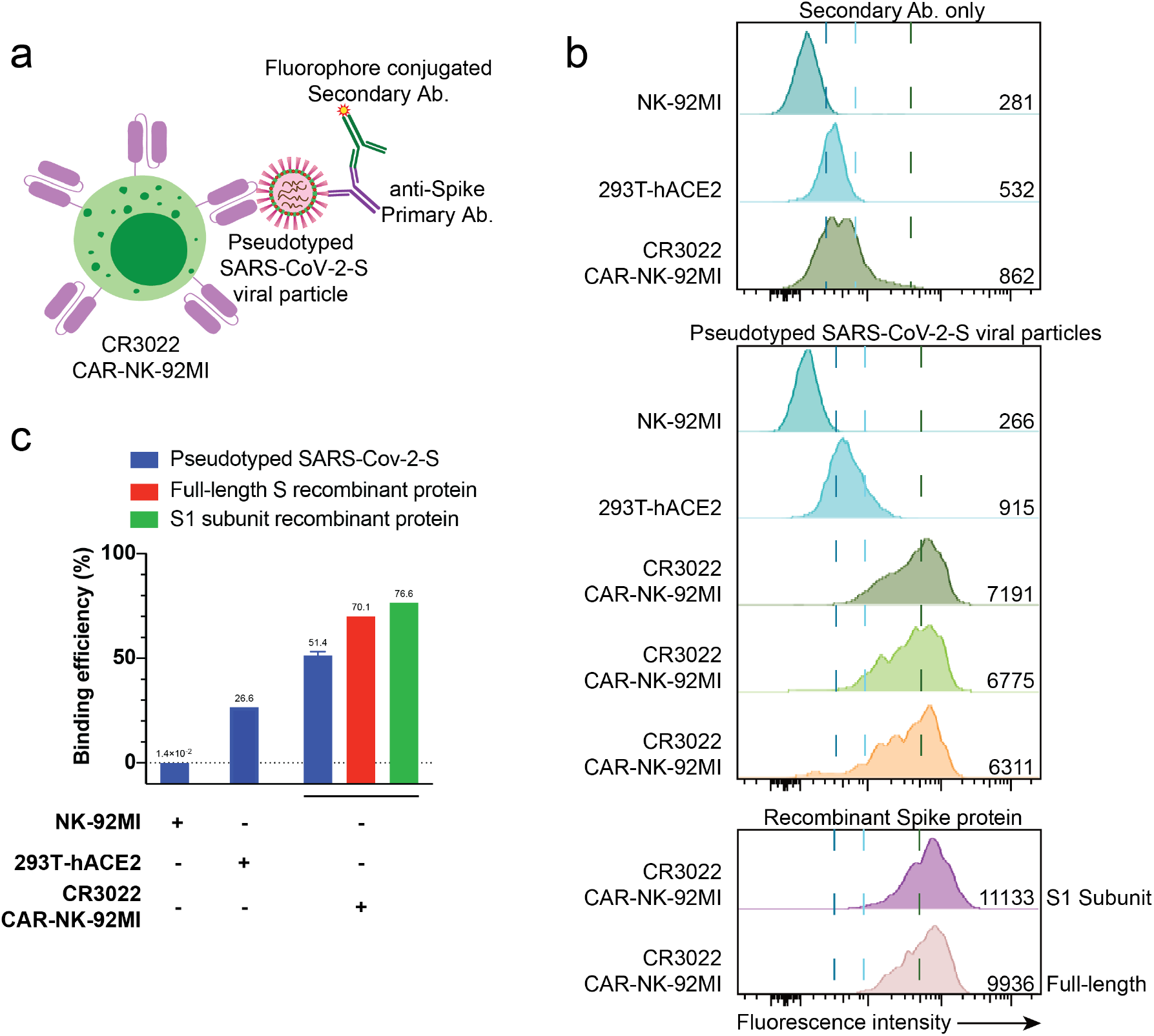
CR3022-CAR-NK-92MI cells bind to pseudotyped SARS-CoV-2-S viral particles. **(a)** Diagram of CR3022-CAR binding to pseudotyped SARS-CoV-2-S viral particles. CR3022-CAR binds to pseudotyped SARS-CoV-2-S viral particle which is then recognized by anti-spike and its corresponding secondary antibody conjugated to a fluorophore. **(b)** Representative histogram of CR3022-CAR binding to pseudotyped SARS-CoV-2-S. CR3022-CAR or NK-92MI or 293T-hACE2 cells were incubated with pseudotyped SARS-CoV-2 or full-length spike or S1 subunit recombinant protein. The binding efficiency was determined by flow cytometry. Experimental sample was performed in triplicate with MFI 6759 ± 440 (a.u.). (**c**) Graph showing the binding efficiency of CR3022-CAR to pseudotyped SARS-CoV-2-S. The values were converted from Figure 2b. Experimental sample was performed in triplicate with binding efficiency 51.4 ± 3.34 (%).

However, the binding efficiency is slightly lower than that of full-length spike recombinant protein (highlighted in red in **Fig. 3c**) and S1 subunit containing RBD protein (highlighted in green in **Fig. 3c**) groups. Surprisingly, 293T-hACE2 cells showed a weaker binding efficiency with the pseudotyped SARS-CoV-2 viral particles, compared to that of CR3022-CAR-NK-92MI cells, indicating that the binding activities of CR3022-CAR-NK-92MI is superior to the natural receptor of SARS-CoV-2 virus. In summary, CR3022-CAR-NK-92MI cells can specifically and strongly bind to the pseudotyped SARS-CoV-2-S viral particles.

### CR3022-CAR-NK cells can be activated by SARS-CoV-2 spike protein receptor binding domain expressing infected target cells and specifically kill their susceptible target cells

After successful establishment of CR3022-CAR-NK cells and demonstration of recombinant His-RBD protein and pseudotyped SARS-CoV-2-S viral particle binding, we further evaluated whether CR3022-CAR-NK cells can be activated by SARS-CoV-2-S infected target cells. To test this, we first transfected the receptor binding domain (RBD) of SARS-CoV-2 spike protein into 293T-hACE2 cells (commonly used cell line for studying the SARS-CoV-2 virus) by transfecting these two cells with an RBD encoding plasmid (**Fig. 4a** and **Fig. 5**). Greater than 90% of transfection efficiencies on 293T-hACE2 cell were obtained. The expression of RBD proteins on 293T-hACE2 cells were verified by flow cytometry (**Fig. 4b** and **Fig. 5**). After successful establishment of these 293T-hACE2-RBD target cells, we tested whether CR3022-CAR-NK-92MI cells can be activated by these target cells using the conventional CD107a assay. Expectedly, the surface level expression of CD107a molecules on CR3022-CAR-NK-92MI cells after co-culturing with these 293T-hACE2-RBD target cells were significantly increased, which was measured by both the percentage and mean fluorescence intensity (MFI) of CD107a on CR3022-CAR-NK92-MI cells, compared to cells cocultured with the parent cells alone (**Fig. 4c**). In addition, the production of TNF-alpha and perforin were greatly increased on CR3022-CAR-NK-92MI cells after co-culturing with these 293T-hACE2-RBD target cells (data not shown).

**Figure 4:**
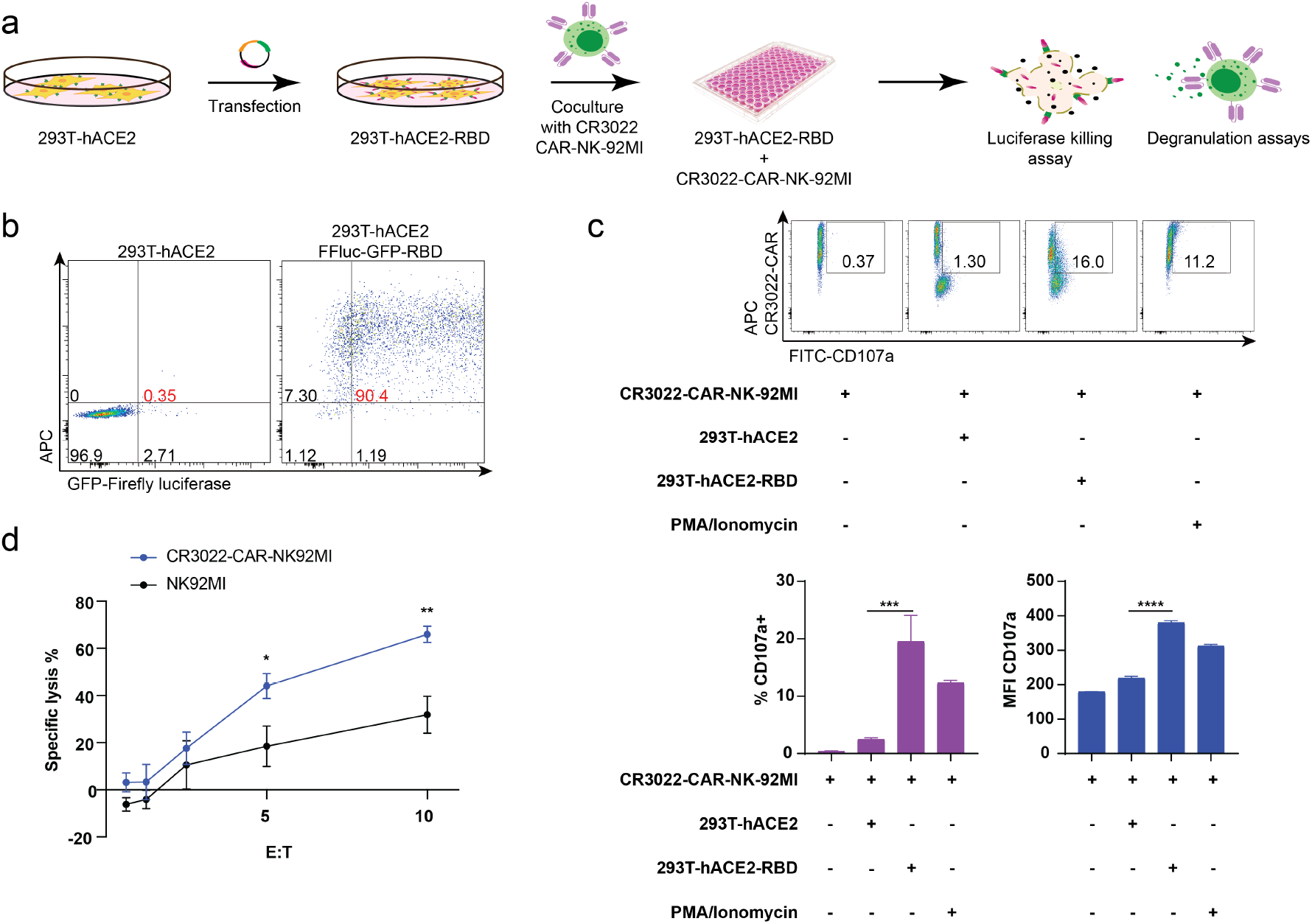
Increased CD107a degranulation and killing activity of CR3022-CAR-NK-92MI cells against 293T-hACE2 cells transfected with RBD-SARS-Cov-2 Spike. (**a**) 293T-hACE2 cells were transfected with a plasmid containing firefly luciferase and GFP as well as SARS-CoV-2 Spike protein receptor binding domain for 48 hours. (**b**) Successful transfection was confirmed by flow cytometry using anti-RBD antibody. Cells were then harvested and used as target cells for subsequent CD107a degranulation assay and luciferase killing assays. (**c**) Representative dot plots of CD107a assay and quantitative data of the percentage and mean fluorescence intensity of CD107a positive CR3022-CAR-NK92MI cells are shown. (**d**) Quantitative data of the luciferase killing assay using CR3022-CAR-NK92MI and wild-type NK-92MI cells against 293T-hACE2-FFLuc-GFP-RBD cells is shown. Experimental groups were performed in triplicate. * p < 0.05, ** p < 0.01, *** p = 0.001, **** p<0.0001 ns p > 0.05. Data represent the mean ± SEM from at least two independent experiments. Briefly, 5 × 10^4^ CR3022-CAR-NK92MI cells were cocultured with either 1 × 10^5^ RBD transfected-293T-hACE2 cells, 293T-hACE2 cells, stimulated with PMA/Ionomycin, or incubated alone for 2 hours at 37°C. Then, cells were harvested, stained for CAR F(ab)2 domain [IgG (H+L)], and CD107a. Representative flow cytometry dot plots, CD107a percent of total CAR cells, and CD107a MFI are shown.

**Figure 5:**
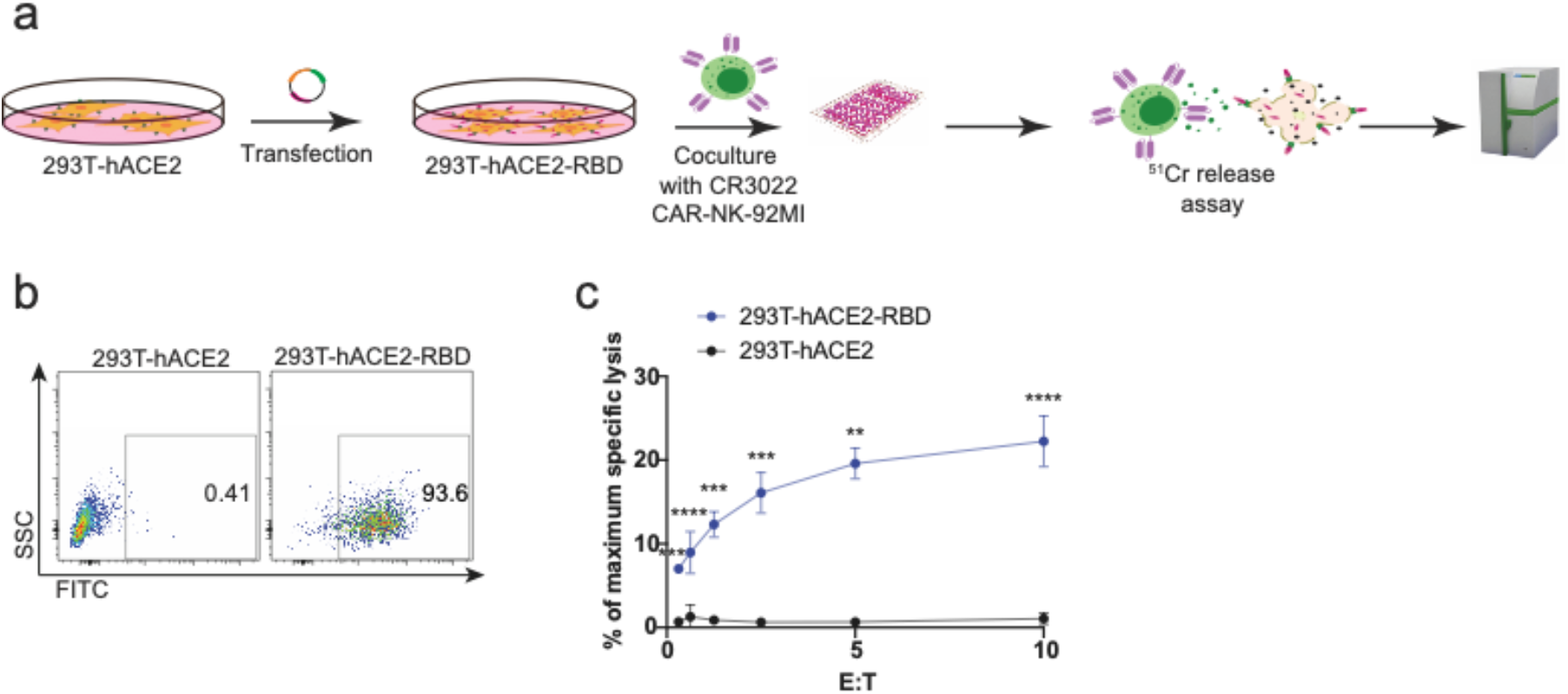
Increased killing activity of CR3022-CAR-NK-92MI cells against 293T-hACE2 cells transfected with SARS-Cov-2 Spike protein Receptor Binding Domain using the ^51^Cr release platform. (a) 293T-hACE2 cells were transfected with SARS-Cov-2 Spike protein receptor binding domain for 48 hours. (b) Successful transfection was confirmed by flow cytometry using Anti-RBD antibody. Cells were then harvested and used as target cells for the subsequent 51Cr release assay. (c) Quantitative data of the ^51^Cr release assay using CR3022-CAR-NK-92MI cells and wild-type NK-92MI cells. Experimental groups were performed in triplicate. * p < 0.05, ** p < 0.01, *** p = 0.001, **** p<0.0001 ns p > 0.05. Data represent the mean ± SEM from at least two independent experiments.

Next, we evaluated the capacity of CR3022-CAR-NK-92MI cells to eradicate SARS-CoV-2 infected target cells (including 293T-hACE2). For 293T-hACE2 cells, an additional plasmid encoding firefly luciferase (FFLuc) tagged with green fluorescent protein (GFP) was also transfected to perform the standard Luciferase assay^2^. Successful luciferase expression on 293T-hACE2 cells was confirmed by flow cytometry analysis (**Fig. 4b**). Compared to control NK-92MI cells, as described previously^17^, CR3022-CAR-NK-92MI cells showed significantly higher killing capacities against 293T-hACE2 cells transfected with RBD (**Fig. 4d**).

To directly test whether CR3022-CAR-NK-92MI cells can kill SARS-CoV-2 infected target cells *in vitro,* we used the 4-hour Chromium-51 (^51^Cr) release assay (a gold standard assay). The data show that CR3022-CAR-NK-92MI cells effectively kill 293T-hACE2-RBD cells by *in vitro* ^51^Cr release assay (**Fig. 5c)**. Thus, by using both the luciferase killing assay platform and ^51^Cr release assay platforms using two different transfected cell lines, we demonstrated the CR3022-CAR-NK-92MI cells can kill the SARS-CoV-2 infected target cells, which supports the clinical use for treatment of SARS-CoV-2 infection.

## DISCUSSION

Recent clinical trials testing cancer immunotherapies have shown promising results for treating infectious diseases^14^. One of crucial barriers to using primary NK cells for immunotherapy is the difficulty in obtaining an adequate number of NK cells from peripheral blood or cold blood before expansion^18^. We have optimized the NK cell expansion technology to buffer this potential limitation. Thus, in this study, we focused on CR3022-CAR-NK-92MI, a NK-92 cell line expressing IL-2 molecule to sustain the persistence *in vivo*^20^.

In this study, we provide proof-of-concept for using CR3022-CAR-based cell therapy for treating severe COVID-19 patients. These experiments will expedite preclinical studies and a potential clinical application during the COVID-19 pandemic.

Although these findings support the therapeutic potential of CR3022-CAR-NK cells for treating severe COVID-19 patients, there are several limitations presented in the current form of study. First, we use the NK-92 cell line in this study. NK-92-mediated immunotherapy is currently undergoing phase I/II clinical trials^21,22^. However, NK-92 cells must be irradiated prior to infusion to prevent permanent engraftment because of malignant potential of NK-92 cells. Second, we use pseudotyped SARS-CoV-2-S viral particles, which is different from the natural SARS-CoV-2 virus. Future studies using natural SARS-CoV-2 virus in the ACE2-transgenic mouse model are needed to test the efficacy and toxicity of CR3022-CAR-NK cells.

In conclusion, development of this novel CAR-NK cell therapy for the treatment of severe COVID-19 patients with maximal efficacy and minimal toxicity will be required to reduce patient risk and enhance the benefit of these expensive and time-intensive therapies. The studies here characterize the biology of CR3022-CAR-NK-92MI cells, test the efficacy of CR3022-CAR-NK-92MI using *in vitro* assays, and finally, define the efficacy of eliminating SARS-CoV-2 infected target cells by CR3022-CAR-NK-92MI cells. This work pioneers the use of CR3022-CAR-NK cells to treat SARS-CoV-2 infected patients and will lead to the development of novel immunotherapeutic strategies for patients presenting with severe COVID-19, and combined with other broadly neutralizing antibodies, will support the development of a universal, “off-the-shelf” CAR-NK based-immunotherapy for COVID-19.

### Online content

Any methods, additional references, source data, and statements of code and data availability are available online.

## METHODS AND MATERIALS

### Antibodies and Reagents

PE anti-human CD3 antibody (clone OKT3), FITC and PE/Cy7 anti-human CD56 antibody (clone HCD56, BioLegend), PE anti-human CD69 antibody (clone FN50, BioLegend), PE anti-human CD8a antibody (clone RPA-T8, BioLegend), APC/Fire 750 anti-human CD226 antibody (DNAM-1) (clone 11A8, BioLegend), APC/Fire 750 antihuman KLRG1 (MAFA) antibody (clone SA231A2, BioLegend), BV421 anti-human CD335 (NKp46) antibody (clone 9E2, BioLegend), PE/Cy7 anti-human CD244 (2B4) antibody (clone C1.7, BioLegend), PE anti-human CD152 (CTLA-4) antibody (clone BNI3), APC anti-human CD366 (Tim-3) antibody (clone F38-2E2), PerCP/Cy5.5 antihuman TIGIT (VSTM3) antibody (clone A15153G), FITC anti-human CD223 (LAG-3) antibody (clone 11C3C65, BioLegend), BV510 anti-human CD314 (NKG2D) antibody (clone 1D11), and APC anti-human CD94 (clone DX22, BioLegend) were purchased from BioLegend (San Diego, CA, USA). APC anti-human CD16 antibody (clone 3G8, BD Biosciences), BV711 anti-human CD314 (NKG2D) antibody (clone 1D11, BD Biosciences), and FITC anti-human CD107a antibody (clone H4A3, BD Biosciences) were purchased from BD Biosciences (San Jose, CA, USA). PE anti-human NKG2C/CD159c antibody (clone 134591, R&D Systems) were purchased from R&D Systems. AF647 Goat anti-human IgG(H+L) F(ab’)_2_ fragment antibody was purchased from Jackson ImmunoResearch (West Grove, PA, USA). Anti-SARS-CoV-2 Coronavirus Spike protein (subunit 1) polyclonal antibody was purchased from Invitrogen (Carlsbad, CA, USA). Anti-SARS-CoV-2 Spike RBD rabbit polyclonal antibody was purchased from SinoBiological (Beijing, China). Anti-His mouse monoclonal antibody IgG1 (clone H-3) was purchased from Santa Cruz Biotechnology (Dallas, TX, USA). Alexa Fluor 488 goat anti-rabbit IgG (H+L) and Alexa Fluor 488 goat anti-mouse IgG1 (g1) were purchased from Fisher Scientific (Waltham, MA).

### Cell lines

293T cell line was purchased from the American Type Culture Collection (ATCC). 293T-hACE2 cell line is a gift from Dr. Abraham Pinter (Rutgers-New Jersey Medical School, PHRI). To maintain the stable expression of hACE2, 293T-hACE2 cells were cultured in DMEM (Corning) supplemented with 10% (v/v) fetal bovine serum (FBS), 100 U/mL Penicillin-Streptomycin (Corning), and 1μg/mL of puromycin at 37°C under 5% (v/v) CO_2_. To establish transient 293T-hACE2-RBD, 293T-hACE2 cells were transfected with 0.5 μg of SARS-CoV-2-RBD plasmid (a gift from Dr. Abraham Pinter) each well in a 24-well plate (Eppendorf) for 48 hours at 37°C under 5% (v/v) CO_2_. Similarly, 293T-hACE2-FFLuc-GFP-RBD cells were transfected with 0.25 μg of SARS-CoV-2-RBD plasmid and 0.25 μg of pHAGE-FFLuc-GFP each well in a 24-well plate (Eppendorf) for 48 hours at 37°C under 5% (v/v) CO_2_. Cells were harvested and immediately used for CD107a degranulation, ^51^Cr release, and FFLuc reporter assays.

### CR3022-CAR construction and retrovirus production

A codon-optimized DNA fragment was synthesized by GENEWIZ encoding the CR3022-specific scFv and sub-cloned into the SFG retroviral vector retroviral backbone in-frame with the hinge component of human IgG1, CD28 trans-membrane domain, intracellular domain CD28 and 4-1BB, and the ζ chain of the human TCR/CD3 complex. The method was previously described^2^, briefly, to produce CR3022-CAR retrovirus, 293T cells were transfected with CR3022-CAR in SFG backbone, RDF, and PegPam3. CR3022-CAR retrovirus was harvested after 48-72 hours and transduced to NK92MI cells in a 24-well plate coated with 0.5 μg/ml of RetroNectin diluted in PBS (Clontech). Two days later, cells were transferred to 75 cm^2^ flask (Corning) and maintained in 35 ml complete NK92MI medium (MEM-a with 12.5% (v/v) FBS, 12.5% (v/v) heat inactivated horse serum, 11 μM ßME, 2 μM folic acid, and 20 μM inositol) supplemented 200 U/mL IL-2 (PeproTech). To determine the expression of CAR, cells were stained for CD56 and anti-human IgG(H+L) F(ab’)2 fragment and analyzed by flow cytometry.

### CR3022-CAR and RBD binding assay

To evaluate the binding activity of CR3022-CAR to RBD domain of SARS-CoV-2-S, CR3022-CAR and NK92MI (5 × 10^5^) cells were incubated with 5 μg of His-gp70-RBD recombinant protein is a gift from Dr. Abraham Pinter in DPBS buffer (0.5 mM MgCl_2_ and 0.9 mM CaCl_2_ in PBS) in for 30 minutes on ice. Cells were washed twice with PBS, stained with anti-His in FACS buffer (0.2% FBS in PBS) for 30 minutes on ice and washed twice with PBS. Cells were then stained with anti-mouse (IgG1) secondary antibody in FACS buffer for 30 minutes on ice, washed twice with PBS, and analyzed by Flow Cytometry.

### CR3022-CAR and pseudotyped SARS-CoV-2-S viral particles binding assay

CR3022-CAR, NK92MI, and 293T-hACE2 (5 × 10^5^) cells were first equilibrated with BM (complete RPMI-1640 containing 0.2% BSA and 10 mM HEPES pH 7.4). Due to the non-specific binding to our CR3022-CAR of our secondary antibody, cells were first blocked with anti-human IgG(H+L) F(ab’)2 fragment for 30 minutes on ice in BM and washed thrice with PBS. Pseudotyped SARS-CoV-2-S (Genscript), full-length recombinant S protein (Acrobio systems), and S1 subunit recombinant protein (a gift from Dr. Abraham Pinter) were diluted with BM to appropriate concentrations. 4 × 10^6^ IFU of pseudotyped SARS-CoV-2-S, or 2 μg of full-length recombinant S protein, or 2 μg of S1 subunit recombinant protein was added to designated wells of a 96-well V bottom plate. Plate was spun at 600 × g for 30 minutes at 32°C, then was incubated at 37°C under 5% (v/v) CO_2_ for 1 hour. Cells were washed twice with PBS, stained with anti-S1 in FACS buffer (0.2% FBS in PBS) for 30 minutes on ice and washed thrice with PBS. Cells were then stained with goat anti-rabbit secondary antibody in FACS buffer for 30 minutes on ice, washed thrice with PBS, and analyzed by Flow Cytometry.

### Flow Cytometry Analysis

NK92MI and CR3022-CAR cells were stained were stained and washed as previously described. Cells were analyzed on a FACS LSRII or an LSR Fortessa flow cytometer (BD). PMT voltages were adjusted and compensation values were calculated before data collection. Data were acquired using FACS Diva software (BD) and analyzed using FlowJo software (BD).

### CD107a Degranulation assay

The CD107a degranulation assay was described previously^3^. Briefly, expanded NK cells (5 × 10^4^) were incubated with 1 × 10^5^ 293T or cells in V-bottomed 96-well plates in complete RPMI-1640 media at 37°C for 2 hours. The cells were harvested, washed, and stained for CD3, CD56, and CD107a with GolgiStop (BD Biosciences) for 30 minutes, and analyzed by flow cytometry.

### FFLuc reporter assay

To quantify the cytotoxicity of CAR-modified immune cells, we also developed the FFLuc reporter system assay. Briefly, an optical 96-well plate (Greiner Bio-One™ No: 655098) was precoated with Retronectin (0.5 μg/ml in PBS) and placed at 4°C overnight. Then, the following day, the wells were aspirated and 293T-hACE2-FFLuc-GFP-RBD and 293T-hACE2 cells were pre-seeded at 1 × 10^4^ target cells/well in 100 μL/well of DMEM supplemented with 10% FBS. The plate was centrifuged for 5 minutes at 350 × g. In a separate 96-well plate, CR3022-CAR-NK-92MI and NK-92MI cells were resuspended at a concentration of 1 × 10^6^ cells/ml. Serial dilutions of effector cells were then prepared according to the effector/target ratio using NK-92MI medium. Then, the effector cells were added to each well of the optical plate (100 μL/well) and incubated at 37°C under 5% (v/v) CO_2_ for 4 hours and then the supernatant was gently discarded. 100 μL of working concentration D-Luciferin was added to each well with the lights turned off. A microplate reader (BioTek, USA) was used to quantify the data. The data was quantified by converting the obtained values to percentage of specific lysis by the following equation: Specific Lysis Percentage (%) = [1-(S-E)/(T-M)]×100, where S is the value of luminescence of the sample well, E is the value of luminescence of the “effector cell only” well compared to the sample well, T is the mean value of luminescence of “Target cell only” wells, and M is the mean value of luminescence of “blank medium only” wells.

### ^51^Cr release assay

To evaluate the cytotoxic activity of CAR-NK cell, the standard 4-hour ^51^Cr release assay was used. Briefly, target cells were labeled with ^51^Cr at 37°C for 2 hours and then resuspended at 1 × 10^5^/mL in NK-92MI culture medium with 10% FBS without IL-2. Then, 1 × 10^4^ target cells were incubated with serially diluted CAR-NK or NK-92MI cells at 37°C for 4 hours. After centrifugation, the supernatants were collected and the released ^51^Cr was measured with a gamma counter (Wallac, Turku, Finland). The cytotoxicity (as a percentage) was calculated as follows: [(sample – spontaneous release) / (maximum release – spontaneous release)] × 100.

### Statistical Analysis

Data were represented as means ± SEM. The statistical significance was determined using a two-tailed unpaired Student t test, a two-tailed paired Student t test, a two-way ANOVA, where indicated. P < 0.05 was considered statistically significant.

## Acknowledgements

We would like to thank the members of the Liu laboratory for their comments on the manuscripts. We thank Dr. Rongfu Wang (Houston Methodist Research Institute) for providing pHAGE-FFLuc-GFP plasmid. We also thank Dr. Abraham Pinter (Rutgers-Public Health Research Institute) for providing 293T-ACE2 cell line. We also would like to thank Dr. Gianpietro Dotti (UNC) for the SFG vectors. This work was supported in part from HL125018 (D. Liu), AI124769 (D. Liu), AI129594 (D. Liu), AI130197 (D. Liu), and Rutgers University-New Jersey Medical School Liu Laboratory Startup funding.

## Author contributions

M.M., S.B., and D.L. designed the study and wrote the manuscript, K.G., assisted with experiments. D.L. supervised the study.

## Competing interests

The authors declare no competing interests.

## Additional information

Supplementary information is available for this paper on the Journal website. Correspondence and requests for materials should be addressed to D.L.

## Notes

### Competing Interest Statement

The authors have declared no competing interest.

## REFERENCE

1. Li, R., et al. Substantial undocumented infection facilitates the rapid dissemination of novel coronavirus (SARS-CoV-2). Science 368, 489–493 (2020).

2. Amanat, F. & Krammer, F. SARS-CoV-2 Vaccines: Status Report. Immunity 52, 583–589 (2020).

3. Ford, N., et al. Systematic review of the efficacy and safety of antiretroviral drugs against SARS, MERS or COVID-19: initial assessment. J Int AIDS Soc 23, e25489 (2020).

4. Sarzi-Puttini, P., et al. COVID-19, cytokines and immunosuppression: what can we learn from severe acute respiratory syndrome? Clin Exp Rheumatol 38, 337–342 (2020).

5. Bloch, E.M., et al. Deployment of convalescent plasma for the prevention and treatment of COVID-19. J Clin Invest 130, 2757–2765 (2020).

6. Xu, X., Ong, Y.K. & Wang, Y. Role of adjunctive treatment strategies in COVID- 19 and a review of international and national clinical guidelines. Mil Med Res 7, 22 (2020).

7. Jean, S.S., Lee, P.I. & Hsueh, P.R. Treatment options for COVID-19: The reality and challenges. J Microbiol Immunol Infect 53, 436–443 (2020).

8. Shah, S., Das, S., Jain, A., Misra, D.P. & Negi, V.S. A systematic review of the prophylactic role of chloroquine and hydroxychloroquine in coronavirus disease- 19 (COVID-19). Int J Rheum Dis 23, 613–619 (2020).

9. Chowdhury, M.S., Rathod, J. & Gernsheimer, J. A Rapid Systematic Review of Clinical Trials Utilizing Chloroquine and Hydroxychloroquine as a Treatment for COVID-19. Acad Emerg Med 27, 493–504 (2020).

10. Dong, L., Hu, S. & Gao, J. Discovering drugs to treat coronavirus disease 2019 (COVID-19). Drug Discov Ther 14, 58–60 (2020).

11. Russell, B., et al. Associations between immune-suppressive and stimulating drugs and novel COVID-19-a systematic review of current evidence. Ecancermedicalscience 14, 1022 (2020).

12. Hu, Y., Tian, Z.G. & Zhang, C. Chimeric antigen receptor (CAR)-transduced natural killer cells in tumor immunotherapy. Acta Pharmacol Sin 39, 167–176 (2018).

13. Bjorkstrom, N.K., Ljunggren, H.G. & Michaelsson, J. Emerging insights into natural killer cells in human peripheral tissues. Nat Rev Immunol 16, 310–320 (2016).

14. Liu, D., et al. Chimeric antigen receptor (CAR)-modified natural killer cell-based immunotherapy and immunological synapse formation in cancer and HIV. Protein Cell 8, 861–877 (2017).

15. Zhou, P., et al. A pneumonia outbreak associated with a new coronavirus of probable bat origin. Nature 579, 270–273 (2020).

16. Tian, X., et al. Potent binding of 2019 novel coronavirus spike protein by a SARS coronavirus-specific human monoclonal antibody. Emerg Microbes Infect 9, 382–385 (2020).

17. Xiong, W., et al. Immunological Synapse Predicts Effectiveness of Chimeric Antigen Receptor Cells. Mol Ther 26, 963–975 (2018).

18. Yan Yang, S.B., Hsiang-chi Tseng, Minh Ma, Ting Liu, Jie-Gen Jiang, Chen Liu, Dongfang Liu. Superior Expansion and Cytotoxicity of Human Primary NK and CAR-NK Cells from Various Sources via Enriched Metabolic Pathways. Molecular Therapy: Methods & Clinical Development in press (2020).

19. Lan, J., et al. Structure of the SARS-CoV-2 spike receptor-binding domain bound to the ACE2 receptor. Nature 581, 215–220 (2020).

20. Tam, Y.K., et al. Characterization of genetically altered, interleukin 2- independent natural killer cell lines suitable for adoptive cellular immunotherapy. Hum Gene Ther 10, 1359–1373 (1999).

21. Arai, S., et al. Infusion of the allogeneic cell line NK-92 in patients with advanced renal cell cancer or melanoma: a phase I trial. Cytotherapy 10, 625–632 (2008).

22. Tonn, T., et al. Treatment of patients with advanced cancer with the natural killer cell line NK-92. Cytotherapy 15, 1563–1570 (2013).

